# Antibody potency, effector function and combinations in protection from SARS-CoV-2 infection *in vivo*

**DOI:** 10.1101/2020.09.15.298067

**Authors:** Alexandra Schäfer, Frauke Muecksch, Julio C. C. Lorenzi, Sarah R. Leist, Melissa Cipolla, Stylianos Bournazos, Fabian Schmidt, Anna Gazumyan, Ralph S. Baric, Davide F. Robbiani, Theodora Hatziioannou, Jeffrey V. Ravetch, Paul D. Bieniasz, Michel C. Nussenzweig, Timothy P. Sheahan

## Abstract

SARS-CoV-2, the causative agent of COVID-19, is responsible for over 24 million infections and 800,000 deaths since its emergence in December 2019. There are few therapeutic options and no approved vaccines. Here we examine the properties of highly potent human monoclonal antibodies (hu-mAbs) in a mouse adapted model of SARS-CoV-2 infection (SARS-CoV-2 MA). *In vitro* antibody neutralization potency did not uniformly correlate with *in vivo* activity, and some hu-mAbs were more potent in combination *in vivo*. Analysis of antibody Fc regions revealed that binding to activating Fc receptors is essential for optimal protection against SARS-CoV-2 MA. The data indicate that hu-mAb protective activity is dependent on intact effector function and that *in vivo* testing is required to establish optimal hu-mAb combinations for COVID-19 prevention.

## Introduction

Coronaviruses (CoV) have a penchant for host range expansion jumping from reservoir species to different hosts resulting in newly emerging human infectious diseases. Indeed, in the past 20 years, three novel human CoV have emerged causing epidemic and pandemic diseases most recently exemplified by SARS-CoV-2, the causative agent of COVID-19 (de Wit et al., 2016, Zhou et al., 2020).

Effective therapeutics are desperately needed to address the COVID-19 pandemic as there are currently no FDA approved therapies and only two treatments authorized for emergency use (remdesivir, convalescent plasma) (U.S. Food & Drug Administration (FDA), 2020). Human monoclonal antibodies (hu-mAbs) hold great potential for treatment and prevention of COVID-19 disease and several potent SARS-CoV-2-specific mAbs targeting multiple non-overlapping epitopes in the receptor binding domain (RBD) in the spike (S) protein have been reported (Robbiani et al., 2020, Baum et al., 2020, Cao et al., 2020, Hansen et al., 2020, Ju et al., 2020, Liu et al., 2020, Pinto et al., 2020, Wang et al., 2020, Zost et al., 2020a, Li et al., 2020). Some of these hu-mAbs have been tested for their ability to prevent or treat SARS-CoV-2 infection in rhesus macaques and hamsters with variable but encouraging results (Rogers et al., 2020, Liu et al., 2020, Shi et al., 2020, Hansen et al., 2020). However, the role of antibody effector function, relative neutralization potency, and combinations in protection have not been examined to date in part because performing experiments in macaques and hamsters under BSL3 conditions is challenging. In addition to the traditional antibody Fc effector functions (i.e. antibody dependent cellular cytotoxicity, phagocytosis etc.), Fc and cellular Fc-receptor interactions drive aspects of both innate and adaptive immunity including macrophage polarization, antigen presentation, and B cell activation. Thus, the Fc-mediated effector functions of neutralizing antibodies may also play a role in shaping diverse aspects of the adaptive immune response.

Small animal models of SARS-CoV-2 replication and pathogenesis are essential for the preclinical development of vaccines and therapeutics. However, SARS-CoV-2 cannot infect standard laboratory mice due to incompatibility between the RBD and the murine ortholog of the human viral entry receptor, angiotensin converting enzyme receptor-2 (mACE2) (Zhou et al., 2020, Walls et al., 2020, Letko et al., 2020). To obviate this problem, we developed an immune competent mouse model of COVID-19 by remodeling the SARS-CoV-2 spike (S) RBD at the mACE2 binding interface (Dinnon et al., 2020). The recombinant virus, SARS-CoV-2 MA, replicates to high titers in the lungs of laboratory mice and has been used to evaluate COVID-19 vaccines and therapeutics including hu-mAbs (Dinnon et al., 2020, Zost et al., 2020a, Corbett et al., 2020). Here, we examine the role of antibody potency, effector function and antibody combinations on protection from SARS-CoV-2 MA infection *in vivo*.

## Results

### Mouse adapting spike mutations have little impact on antibody neutralization in vitro

To determine if the two amino acid changes in the SARS-CoV-2 MA RBD (Q498T/P499Y) would affect antibody neutralization we used an HIV-1 virus pseudotyped with the SARS-CoV-2-MA S protein (S-MA). The recombinant pseudovirus was produced by co-transfection of S-MA with a replication incompetent proviral genome (NL4-3ΔEnv-NanoLuc) which lacks a functional env gene and encodes a nanoluciferase reporter in the place of the nef gene (Fig. 1A) (Robbiani et al., 2020, Schmidt et al., 2020). To maximize S-incorporation, we truncated the C-terminus of S-MA by 19 amino acids. Using nanoluciferase reporter expression as a measure of infection, we compared the ability of pseudotyped virus bearing either WT SARS-CoV-2 S (wtS) or S-MA to infect HT1080 cells stably expressing either human or murine ACE2. Consistent with previous reports (Zhou et al., 2020, Dinnon et al., 2020), wtS did not support pseudovirus infection of murine ACE2 expressing cells (Fig. 1B). In contrast, the mouse adapted SARS-CoV-2 supported robust infection of both mouse and human ACE2 expressing cells (Fig. 1B).

**Figure 1:**
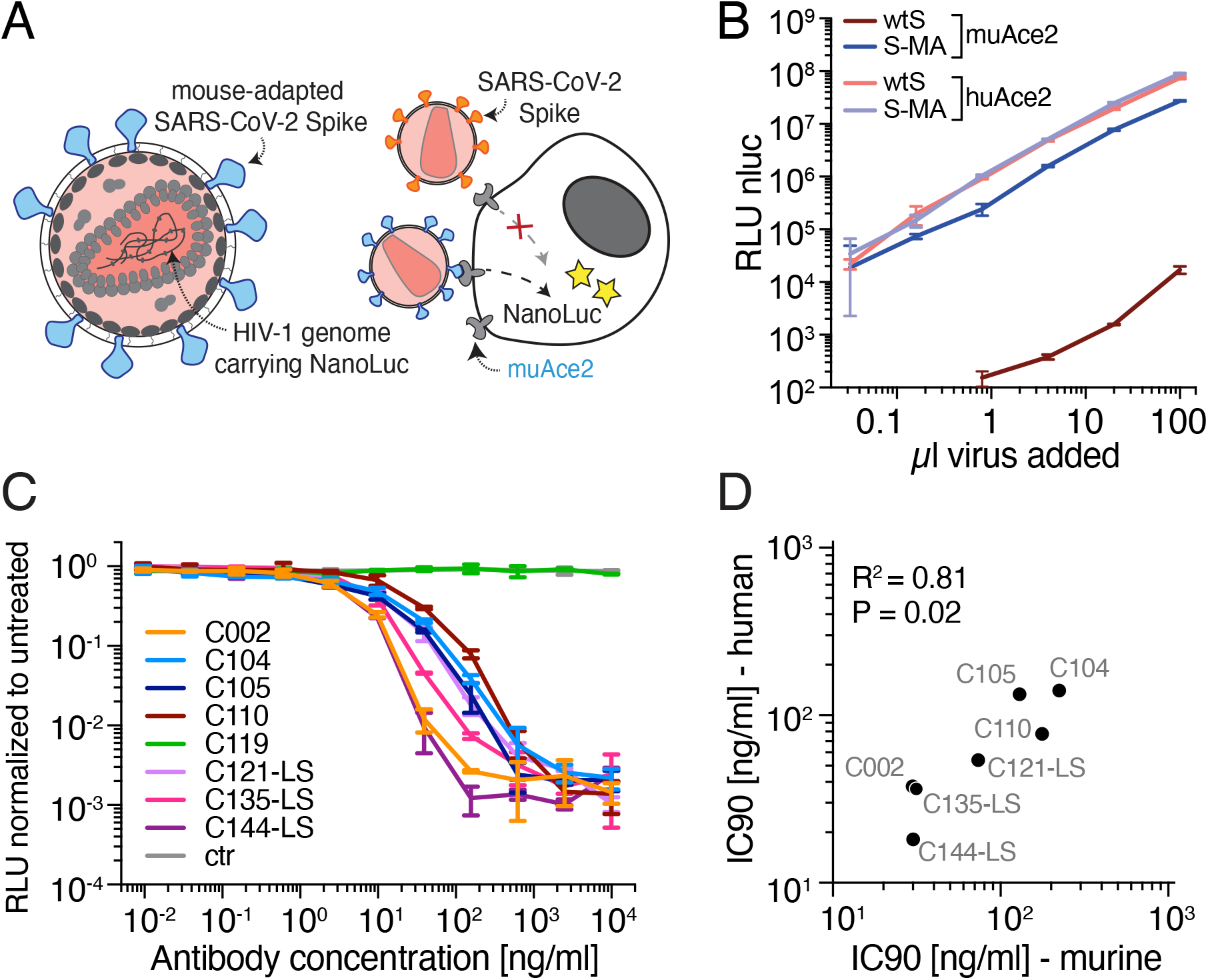
Antibody potency against the mouse adapted SARS-CoV-2 spike. **A.** Diagram of the mouse-adapted (MA) SARS-CoV-2 pseudovirus luciferase assay. SARS-CoV-2 mouse-adapted spike (SARS-CoV-2 S-MA) pseudotyped HIV-1 particles carrying the nanoluc gene are used to infect murine (mu) Ace2-expressing HT1080 cells, which will express nanoluc luciferase upon infection, while SARS-CoV-2 spike (wtS) pseudotyped particles are unable to infect muAce-expressing cells. **B.** Relative luminescence unit (RLU) reads from lysates of muAce2 and human (hu) Ace2-expressing HT1080 cells infected with increasing amounts of SARS-CoV-2 S-MA and wtS pseudovirus. Data are mean +/− s.d. of triplicates. One representative experiment is shown. **C.** The normalized relative luminescence values for cell lysates of HT1080_muAce2_ cells 48h after infection with SARS-CoV-2 S-MA pseudovirus in the presence of increasing concentrations of monoclonal antibodies. n = 8 samples and 1 isotype control. Data are mean +-s.d. of duplicates. One representative experiment is shown. **D.** IC_90_ values detected in the SARS-CoV-2 S-MA pseudovirus neutralization assay (IC_90_-murine) plotted against those detected in the wtS SARS-CoV-2 pseudovirus neutralization assay (IC_90_ - human). r = 0.8095, P < 0.02178. Mean values of at least two experiments are shown.

The S-MA pseudotyped virus was used to measure the neutralizing activity of 8 different IgG1 hu-mAb with variable potencies against SARS-CoV-2 ranging from IC_50_/IC_90_ of 4.4/18 to 26/140 ng/ml (Table 1) (Robbiani et al., 2020). C002, C104, C105, C119, C121, C144 all target the hACE2 interaction surface of the RBD albeit at different angles of approach, and C135 and C110 target a separate non-overlapping epitope within the RBD (Barnes et al., 2020b, Robbiani et al., 2020). With the exceptions of C119, the antibody neutralization titers were similar when assayed using mouse adapted S-MA or wtS pseudotyped viruses (R = 0.81 p = 0.02, Table 1, Fig. 1C and Fig.1D). C119 was inactive against S-MA pseudotyped virus because the epitope targeted by this antibody overlaps the mouse adapting mutations (Fig. 1C) (Dinnon et al., 2020, Barnes et al., 2020a). Overall, however mouse adapting mutations in the S protein do not significantly affect neutralization by most RBD targeting antibodies tested *in vitro*.

**Table 1.** Inhibitory concentrations of monoclonal antibodies.

| Antibody ID | murine |  | human |  |
| --- | --- | --- | --- | --- |
|  | IC50 (ng/ml) | IC90 (ng/ml) | IC50 (ng/ml) | IC90 (ng/ml) |
| C002 | 3.7 | 29.9 | 8.9* | 37.6* |
| C002-GRLR | 3.8 | 22.7 |  |  |
| C104 | 6.8 | 223.9 | 23.3* | 140.3* |
| C104-GRLR | 15.4 | 160.5 |  |  |
| C105 | 8.0 | 129.8 | 26.1* | 133.7* |
| C110 | 22.6 | 176.9 | 18.4* | 77.3* |
| C110-GRLR | 8.1 | 267.5 |  |  |
| C119 | >1000 | >1000 | 9.1* | 97.8* |
| C121-LS | 6.5 | 73.5 | 10.7 | 54.0 |
| C135-LS | 6.9 | 31.5 | 17.3 | 36.3 |
| C144-LS | 3.2 | 30.2 | 4.4 | 18.2 |

\* (Robbiani et al., 2020)

### *In vitro* antibody neutralization does not uniformly correlate with their *in vivo* efficacy

To determine if there is a correlation between *in vitro* neutralization and *in vivo* activity, we performed prophylactic efficacy studies in aged BALB/c mice. Monoclonal antibodies (8 mg/Kg) were administered by intraperitoneal injection 12hr before intranasal infection with 1×10^5^ plaque forming units (PFU) of SARS-CoV-2 MA (Fig. 2A). Virus lung titers were measured by plaque assay two days after infection, which is the kinetic peak of viral replication in this model (Dinnon et al., 2020). Since this is primarily a virus replication model, infected mice did not display overt disease. Mice injected with the isotype control antibody (anti Zika antibody 3633, (Robbiani et al., 2017)) had mean viral lung titers of 1×10^6^ PFU (Fig. 2A, Table 2). In agreement with the *in vitro* neutralization data, C119 failed to protect against SARS-CoV-2 MA *in vivo* (Figs. 1C and 2A, Table 2). In contrast, the other anti-SARS-CoV-2 antibodies tested protected against infection to varying degrees (Fig. 2A). C104 (IC_90_ 223 ng/ml) reduced viral loads in the lungs of all mice to below the limit of detection (i.e. 50 PFU). Other antibodies that were more potent against SARS-CoV-2-MA pseudoviruses than C104 *in vitro* lowered viral loads by 3-4 orders of magnitude (Fig. 2A; C002, C110, C121-LS, C135-LS, and C144-LS). Interestingly, C105 only reduced the viral loads *in vivo* by 1-2 orders of magnitude yet its neutralizing activity against SARS-CoV-2 MA pseudotyped virus *in vitro* was similar to C104 (Figs. 1, 2 and Table 1). Comparison of the mean virus lung titer and respective IC_90_ revealed that the *in vitro* neutralizing activity in pseudovirus assays did not uniformly correlate with *in vivo* efficacy (Fig. 2B).

**Table 2.** SARS-CoV-2 MA virus lung titers with single hu-mAb prophylaxsis.

| Antibody ID | C002 | C104 | C105 | C110 | C119 | C121-LS | C135-LS | C144-LS | isotype control |
| --- | --- | --- | --- | --- | --- | --- | --- | --- | --- |
| Dose Level (mg/Kg) | 8 | 8 | 8 | 8 | 8 | 8 | 8 | 8 | 8 |
| Number of animals | 10 | 10 | 10 | 10 | 9 | 10 | 9 | 10 | 10 |
| Minimum (PFU/mL) | 4.5E+01 | 4.5E+01 | 2.5E+03 | 4.5E+01 | 1.0E+05 | 1.5E+02 | 4.5E+01 | 4.5E+01 | 4.5E+05 |
| Maximum (PFU/mL) | 7.5E+03 | 4.5E+01 | 6.3E+04 | 6.0E+03 | 1.5E+06 | 7.5E+03 | 3.5E+03 | 2.0E+03 | 3.0E+06 |
| Range | 7.5E+03 | 0.0E+00 | 6.0E+04 | 6.0E+03 | 1.4E+06 | 7.4E+03 | 3.5E+03 | 2.0E+03 | 2.6E+06 |
| Geometric mean (PFU/mL) | 9.9E+02 | 4.5E+01 | 1.3E+04 | 2.2E+02 | 4.9E+05 | 1.1E+03 | 3.3E+02 | 1.5E+02 | 1.0E+06 |
| Geometric SD factor | 8.6 | 1.0 | 2.6 | 6.9 | 2.5 | 4.0 | 4.8 | 5.0 | 1.8 |

**Figure 2.**
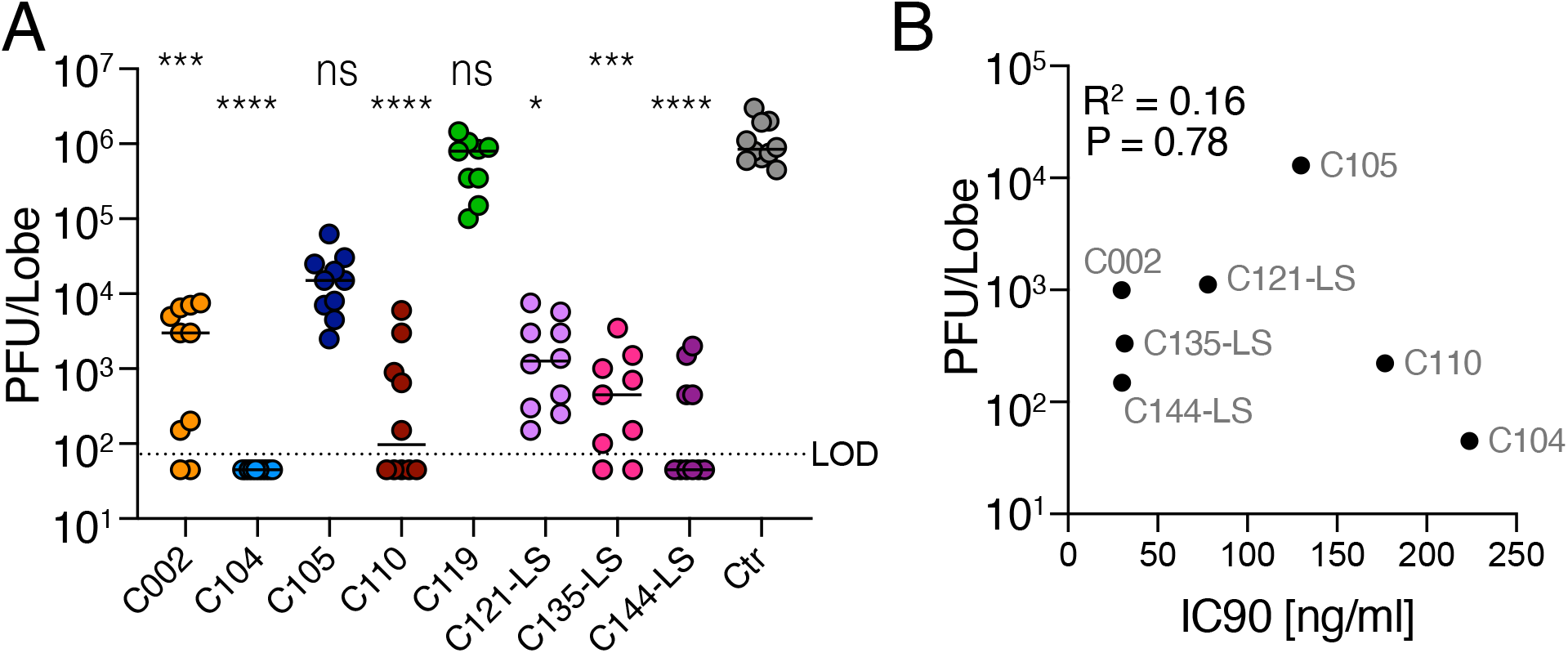
In vivo potency does not uniformly correlate with in vitro potency. **A.** SARS-CoV-2 MA lung titer following antibody prophylaxis. Antibodies (8mg/kg) were delivered intraperitoneally 12hr prior to infection with 1×10^5^ particle forming units (PFU) of SARS-CoV-2 MA. Combined data from two independent experiments is shown. All groups are N = 10 mice/group except for C119 and C135-LS. The line is at the geometric mean and each symbol represents the titer for a single animal. Asterisks indicate statistical differences as compared to isotype control by one-way ANOVA with a Dunn’s multiple comparison test (P < 0.0001, ****; P = 0.003, ***; P = 0.007-0.004, **). **B.** PFU/Lobe values plotted against IC_90_ values detected in the SARS-CoV-2 S-MA pseudovirus neutralization assay. R = 0.1585, P = 0.788. The R and P values in **A.** and **B.** were determined by two-tailed Spearman’s correlations.

### Antibody Fc-effector function mediates protection against SARS-CoV-2 MA *in vivo*

Virus neutralization *in vitro* is independent of antibody Fc effector functions that impact *in vivo* efficacy against other viral infections (Lu et al., 2016, Halper-Stromberg et al., 2014, DiLillo et al., 2014, Bournazos et al., 2019, DiLillo et al., 2016, Bournazos et al., 2014). To examine the role of Fc-effector function on protective activity against SARS-CoV-2 MA *in vivo* we introduced the G236R/L328R (GRLR) mutation that abrogates antibody Fc receptor interaction in C002, C104 and C110 (Fig. 3) (Bournazos et al., 2014). C002 and C104 target epitopes on the ACE2 binding interface of RBD while C110 targets RBD but does not directly overlap the ACE2 interaction surface (Robbiani et al., 2020, Barnes et al., 2020a). As expected, the three antibody variants (C002_GRLR_, C104_GRLR_, C110_GRLR_) had IC_90_ values that were not significantly different from wild type hu-mAb in SARS-CoV-2 MA pseudovirus assays *in vitro* (Fig. 3A and B). Elimination of Fc-effector function did not significantly affect *in vivo* protection by C002, the least potent of the three antibodies (Fig. 3C, Table 3, C002 = 9.9×10^2^ PFU vs. C002-GRLR = 1.6.×10^3^ PFU, P = 0.72). In contrast, loss of Fc-effector function significantly decreased the potency of both C104_GRLR_ and C110_GRLR_ (Fig. 3C). Fc null C104_GRLR_ was 14-fold, and C110_GRLR_ 6-fold less potent than their Fc effector sufficient counterparts (P = 0.0001 and P = 0.004, respectively, Fig. 3C, Table 3). Variable Fc effector requirements were also observed for anti-flu antibodies, suggesting that nature of the antibody-pathogen interaction can influence the ability of the Fc to engage its receptor (DiLillo et al., 2016).

**Table 3.** SARS-CoV-2 MA virus lung titers with single hu-mAb prophylaxsis using GRLR Fc mutatants and murine Fc region grafts.

| Antibody ID | C002 | C002 GRLR | C104 | C104 GRLR | C110 | C110 GRLR | isotype control |
| --- | --- | --- | --- | --- | --- | --- | --- |
| Dose Level (mg/Kg) | 8 | 8 | 8 | 8 | 8 | 8 | 8 |
| Number of animals | 10 | 10 | 10 | 10 | 10 | 10 | 10 |
| Minimum (PFU/mL) | 4.5E+01 | 4.5E+01 | 4.5E+01 | 4.5E+01 | 4.5E+01 | 6.5E+02 | 1.0E+05 |
| Maximum (PFU/mL) | 7.5E+03 | 1.1E+04 | 4.5E+01 | 3.0E+03 | 6.0E+03 | 2.5E+04 | 1.1E+06 |
| Range | 7.5E+03 | 1.1E+04 | 0.0E+00 | 3.0E+03 | 6.0E+03 | 2.4E+04 | 1.0E+06 |
| Geometric mean (PFU/mL) | 9.9E+02 | 1.6E+03 | 4.5E+01 | 6.9E+02 | 2.2E+02 | 3.4E+03 | 4.8E+05 |
| Geometric SD factor | 8.6 | 5.9 | 1.0 | 3.2 | 6.9 | 3.0 | 1.8 |

| Antibody ID | C104 IgG1 | C104 IgG2 | C104 D265A | C004 GRLR | C004 | isotype control |
| --- | --- | --- | --- | --- | --- | --- |
| Dose Level (mg/Kg) | 8 | 8 | 8 | 8 | 8 | 8 |
| Number of animals | 5 | 5 | 5 | 10 | 10 | 5 |
| Minimum (PFU/mL) | 4.5E+01 | 4.5E+01 | 1.5E+03 | 4.5E+01 | 4.5E+01 | 1.2E+06 |
| Maximum (PFU/mL) | 2.0E+04 | 2.0E+02 | 2.5E+04 | 3.0E+03 | 4.5E+01 | 5.5E+06 |
| Range | 2.0E+04 | 1.6E+02 | 2.4E+04 | 3.0E+03 | 0.0E+00 | 4.3E+06 |
| Geometric mean (PFU/mL) | 7.5E+02 | 6.2E+01 | 7.4E+03 | 6.9E+02 | 4.5E+01 | 3.4E+06 |
| Geometric SD factor | 15.4 | 1.9 | 3.8 | 3.2 | 1.0 | 1.8 |

**Figure 3:**
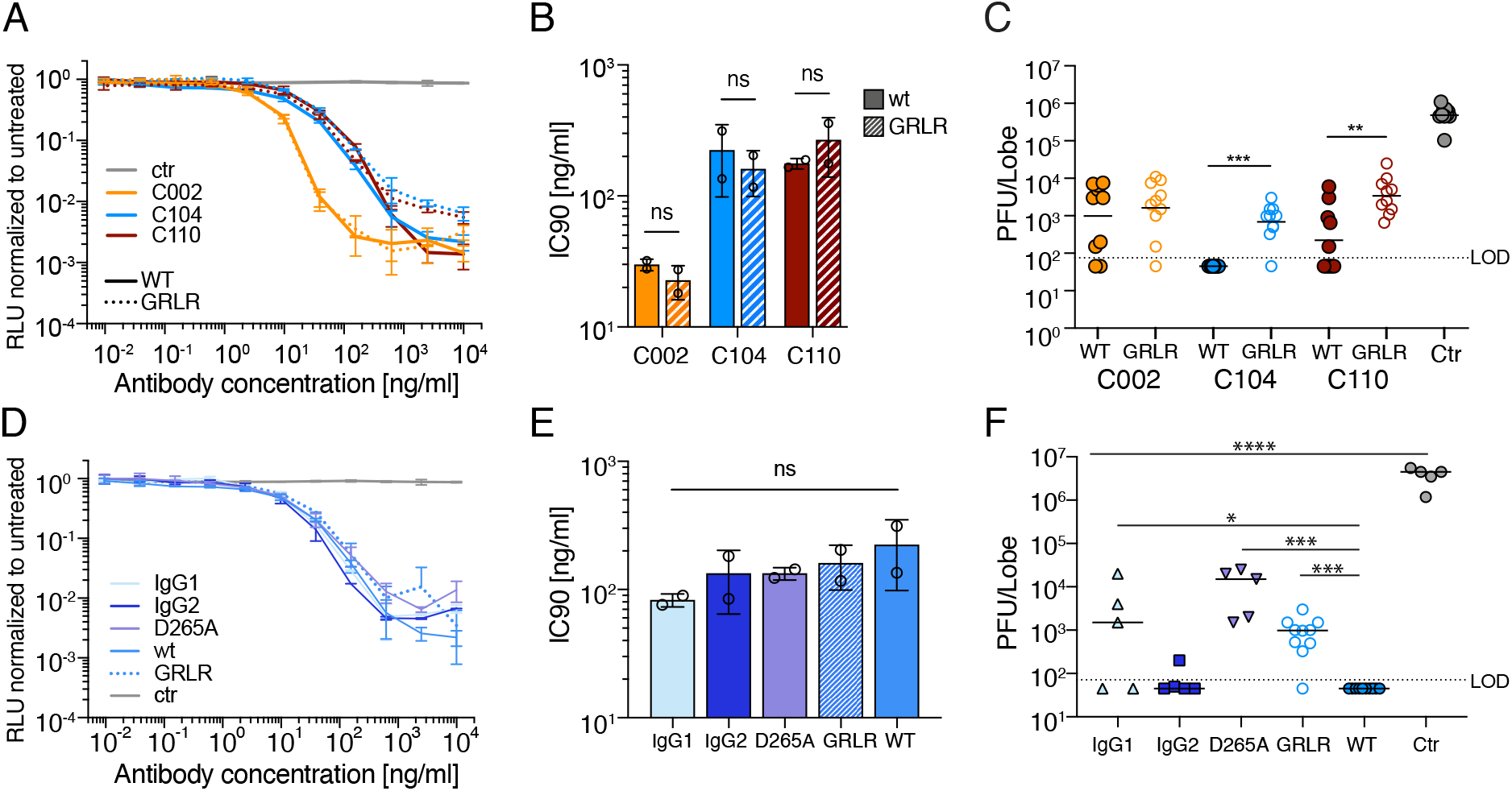
The variable requirement of Fc-effector function for SARS-CoV-2 neutralization. **A.** Antibody potency curves for WT and GRLR mutant antibodies. The normalized relative luminescence values for cell lysates of HT1080_muAce2_ cells 48 h after infection with SARS-CoV-2 S-MA pseudovirus in the presence of increasing concentrations of WT (solid lines) or GRLR-modified monoclonal antibodies (dotted lines). n = 3 samples. Data are mean +/− s.d. of duplicates and one representative experiment is shown. **B.** IC_90_ values for antibodies shown in **A.** Bars represent mean values of two experiments (shown as open circles), error bars indicate standard deviation. **C.** SARS-CoV-2 MA lung titer following antibody prophylaxis. WT or GRLR antibodies (8mg/kg) were delivered intraperitoneally 12hr prior to infection with 1×10^5^ PFU of SARS-CoV-2 MA. Combined data from two independent experiments is shown and all groups are N = 10 mice/group. The line is at the geometric mean and each symbol represents the titer for a single animal. Asterisks indicate statistical differences as compared to isotype control by Mann-Whitney test (P < 0.004, **; P = 0.0001, ***). **D.** Antibody potency curves for WT and Fc mutant antibodies performed and displayed similar to that in panel **A**. **E.** IC_90_ values for the Fc mutant antibodies shown in D. Bars represent mean values of two experiments (shown as open circles), error bars indicate standard deviation. **F.** SARS-CoV-2 MA lung titer following antibody prophylaxis with grafted the variable domains of C104 into mouse IgG1, IgG2b and IgGD265A mAb. WT C104 and isotype control antibody treated groups were controls. Antibodies (8mg/kg) were delivered intraperitoneally 12hr prior to infection with 1×10^5^ PFU of SARS-CoV-2 MA. One independent experiment is shown. All groups are N = 5 mice/group. The line is at the geometric mean and each symbol represents the titer for a single animal. Asterisks (****) indicate statistical differences as compared to isotype control by one-way ANOVA with Dunnet’s multiple comparison test (P < 0.0001) or Mann-Whitney test (P = 0.022, *; P = 0.0001 to 0.0003, ***).

To determine which Fc receptors are responsible for enhanced protective activity we grafted the variable domains of C104 onto the mouse IgG1, IgG2a Fc and the mouse IgG1 Fc variant IgGD265A, which is null binding mutant for all mouse Fc receptors (Clynes et al., 2000). Mouse subclasses display differential affinity for activating (FcRI, III and IV) and the inhibitory (FcRIIB) receptors (Nimmerjahn and Ravetch, 2005). Since activating and inhibitory receptors are co-expressed on most effector cells, the *in vivo* activity of an IgG Fc is the result of the differential affinity of Fc binding to these receptors. Mouse IgG1 binds preferentially to FcRIIB, an inhibitory receptor, while IgG2a binds primarily FcRIV receptor, an activating receptor. FcRIV is expressed on monocytes/macrophages, neutrophils and dendritic cells but not on murine NK cells (Bournazos et al., 2017). As expected, the three mouse antibody variants had neutralization potencies that were not significantly different from each other or their human IgG1 counterpart in SARS-CoV-2 S-MA pseudovirus assays *in vitro* (Fig. 3D and E, Table 3). Whereas mouse C104-IgG2a provided a similar level of protection as WT human C104 (human IgG1, a human subclass that engages mouse FcRIV), mouse C104-IgG1 and the C104-IgGD265A variant were significantly less active (Fig. 3F, Table 3). We conclude that activating Fc receptors are essential for optimal protection against SARS-CoV-2 MA *in vivo*.

### Antibody combinations potently neutralize SARS-CoV-2 in vivo

Combinations of antibodies targeting non-overlapping epitopes have the potential to increase anti-viral potency and can prevent the emergence of antibody escape mutations (Hansen et al., 2020), (Weisblum et al., 2020). To test whether there might be a benefit of antibody combinations for *in vivo* protection, we tested mixtures of antibodies that target non-overlapping epitopes on the RBD (C135/C121 and C135/C144) (Robbiani et al., 2020). The antibody mixtures were tested at combined total doses of 16, or 5.3 or 1.8 mg/Kg compared to 8mg/Kg of each antibody alone or 16mg/Kg of isotype control. As expected, the isotype control antibody failed to reduce virus replication (Fig. 4A). While 16mg/Kg of C135/121 provided sterilizing protection from SARS-CoV-2 MA replication in all mice tested (N = 15), C135/C144 performed nearly as well with only 1 mouse of 15 tested having a low but measurable virus titer (Fig. 4A, Table 4). Similarly, both combinations at 5.3 mg/Kg provided significant protection from replication similar to that of 16 mg/Kg. Whereas the C135/C144 was sterilizing at 5.3 mg/Kg, half of the C135/C121 mice had low but detectable virus lung titers. Notably, the 1.8 mg/Kg dose of either combination significantly reduced titers as compared to isotype control antibody but was not as effective as 5.3 or 16 mg/Kg (Fig. 4A, Table 4). In addition, antibody combinations improved the levels of protection achieved from single antibodies alone (Fig. 4B). These data argue that some antibody combinations can neutralize SARS-CoV-2 more potently than each antibody alone.

**Table 4.**
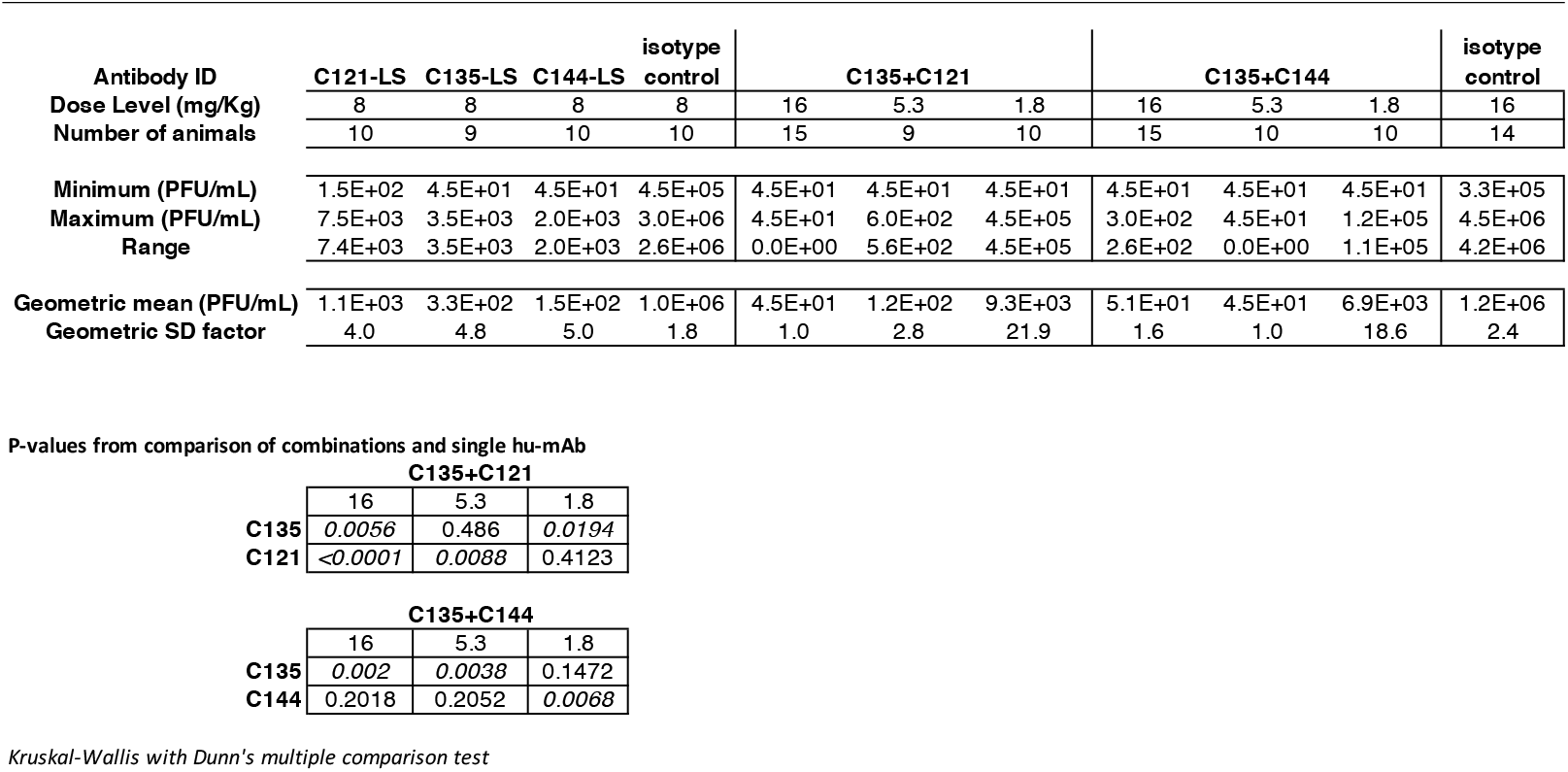
Comparison of SARS-CoV-2 MA virus lung titers with either single hu-mAb or hu-mAb combination prophylaxis.

**Figure 4:**
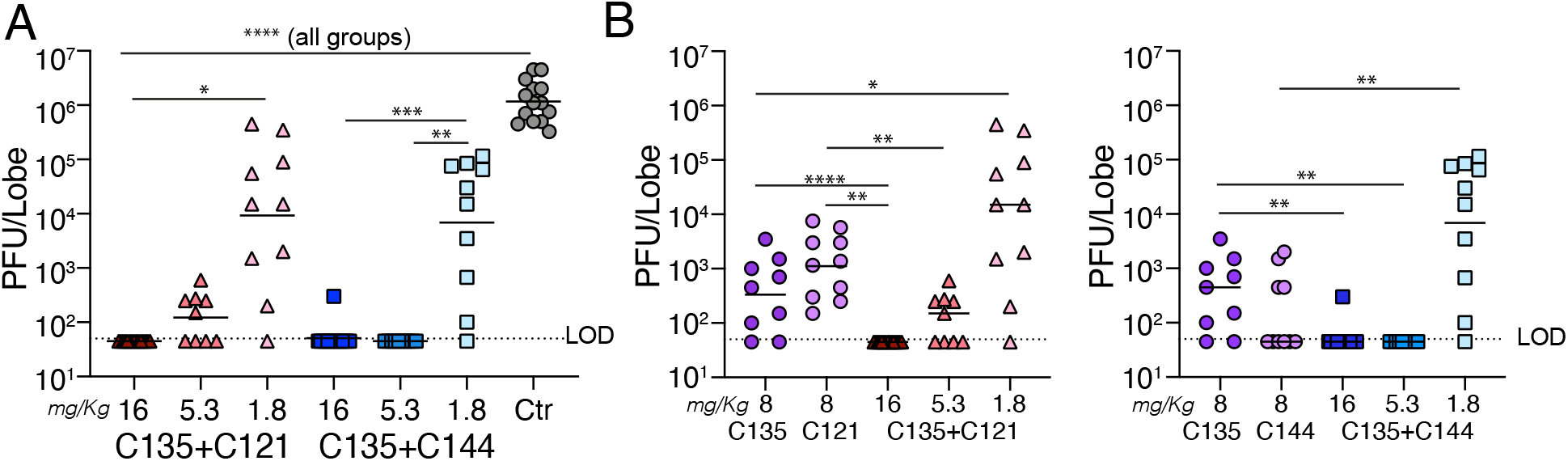
Antibody combinations synergize to increase in vivo potency. **A.** SARS-CoV-2 MA lung titer following antibody prophylaxis with isotype control (16mg/Kg) or combinations of C135+121 or C135+C144 mixed at a ratio of 1:1 for combined dose levels of 16, 5.3 or 1.8 mg/Kg. Antibodies were delivered intraperitoneally 12hr prior to infection with 1×10^5^ PFU) of SARS-CoV-2 MA. Combined data from two independent experiments is shown. For 16mg/kg groups, N = 14-15 mice and all other groups were 9-10 mice/group. The line is at the geometric mean and each symbol represents the titer for a single animal. Asterisks indicate statistical differences as compared to isotype control by one-way ANOVA with a Dunnet’s multiple comparison test (P < 0.0001, ****) or ANOVA with a Tukey’s multiple comparison test (P = 0.0005, ***; P = 0.0015, **; P = 0.02, *). **B.** Comparison of SARS-CoV-2 MA lung titers with single antibodies at 8 mg/Kg described in Fig. 2A or combinations described in “**A**”. Differences were determined by Kruskal-Wallis test with a Dunn’s multiple comparison test (P = <0.0001, ****; P = 0.001-0.008, **; P = 0.01, *).

## Discussion

To date there have been over 26 million cases and over 850,000 deaths attributed to COVID-19 globally, 20% of which have occurred in the United States (Johns Hopkins University (JHU), 2020). In addition to the many vaccines and drugs now being tested, passive transfer of potent hu-mAb holds great promise for COVID-19 prevention and therapy. Neutralizing hu-mAb targeting SARS-CoV-2 promote reduction in viral load and prevent infection in macaques and hamsters (Hansen et al., 2020, Rogers et al., 2020, Ju et al., 2020). Here, we examined the relationship between antibody neutralizing potency in *in vitro* assays and protection in a small animal model of SARS-CoV-2 infection (Dinnon et al., 2020). The results suggest that engagement of activating Fc receptors and that some antibody combinations can enhance the efficacy of anti-SARS-CoV-2 antibodies *in vivo*.

The neutralizing activity of antibodies to SARS-CoV-2 has primarily been tested *in vitro* using pseudotype viruses and microneutralization assays (Schmidt et al., 2020). How these *in vitro* results translate to *in vivo* protective activity had not been determined. Our results indicate that the relationship between neutralizing activity against SARS-CoV-2 MA *in vitro* and antiviral activity *in vivo* is not linear. One potentially important factor is the role of leukocyte Fc receptors on viral clearance and infected cell killing *in vivo* (Bournazos et al., 2020). Indeed, Fc receptors play an important role in viral clearance during HIV-1, Ebola and Influenza infections (Lu et al., 2016, Halper-Stromberg et al., 2014, DiLillo et al., 2014). Our experiments indicate that Fc receptors are also essential for optimal antibody mediated protection against SARS-CoV-2. Among this family of receptors, activating Fc receptors on macrophages, neutrophils and dendritic cells are critical for enhanced antibody protection against SARS-CoV-2 MA (Mercado et al., 2020). Thus far, the evidence from both SARS-CoV-2 vaccine and hu-mAb studies does not support the notion that antibody dependent enhancement (ADE) of infection driven by Fc receptor engagement occurs with SARS-CoV-2 as is observed for flavivirus such as dengue virus (Halstead and Katzelnick, 2020, Laczko et al., 2020). In addition, exacerbation of dengue virus infection by ADE is driven by antibody-dependent increases in infection frequency of antigen presenting cells (monocytes, macrophages, dendritic cells, etc.) normally targeted by dengue virus while the major cellular targets of SARS-CoV-2 are respiratory epithelial cells (Wang et al., 2017). Thus, there is currently little evidence to suggest that passively transferred immunity via hu-mAb therapy will initiate immune pathologies such as ADE.

An additional non-mutually exclusive explanation for the disparity between the *in vivo* antiviral activity and *in vitro* neutralization results relates to the heterogeneity in the way neutralizing antibodies target the SARS-CoV-2 RBD domain (Barnes et al., 2020b, Barnes et al., 2020a). Antibodies can neutralize by binding to the ACE2 interaction surface thereby blocking RBD interaction with its cellular receptor directly or indirectly. Even among the antibodies that directly block the interaction between the RBD and ACE2, there is significant heterogeneity in terms of their angles of approach to binding. Among neutralizing antibodies that target the RBD C144 belongs to a particularly potent class whose mechanism of action involves blocking RBD-ACE-2 interaction and additionally locking the RBD in a closed configuration making it inaccessible to ACE2 (Barnes et al., 2020b, Barnes et al., 2020a). C002, C104 and C105 also bind to the ACE2 interacting surface of the RBD but they approach the RBD from different angles and bind by different mechanisms (Barnes et al., 2020b, Barnes et al., 2020a). C104 is the least potent neutralizer of the three antibodies *in vitro* but the most effective against SARS-CoV-2 MA *in vivo*. Among other factors, the angle of approach of an antibody to the RBD may alter its *in vivo* potency by influencing accessibility of the Fc domain to its receptor on effector cells, as suggested for neutralizing antibodies to influenza (DiLillo et al., 2016).

*In vitro* experiments with chimeric VSV-SARS-CoV-2 viruses indicate that antibodies can select for escape mutants and that combinations of antibodies targeting non-overlapping sites can prevent the emergence of resistant variants (Hansen et al., 2020, Weisblum et al., 2020). Antibody combinations also have the potential to act synergistically, but there is little evidence for synergy *in vitro* (Robbiani et al., 2020, Liu et al., 2020, Rogers et al., 2020, Zost et al., 2020b). Nevertheless, combinations of antibodies were more potent than their individual components in protecting against SARS-CoV-2 MA infection. Antibody combinations that target non-overlapping epitopes may be especially promising for clinical development because they can be dose sparing and also prevent selection of resistant variants.

In summary, the data supports the idea that specific combinations of antibodies with intact Fc effector function and should be developed for optimal protection against SARS-CoV-2.

## Material and Methods

### Cells and viruses

Human Ace2-expressing HT1080 cells (HT1080_Ace2_cl.14) were described previously (Schmidt et al., 2020). For constitutive expression of murine Ace2 (muAce2) in HT1080 cells, a cDNA encoding muAce2 was inserted into a lentiviral vector CSIB 3’ to the SFFV promoter. HT1080_muAce2_ cells were generated by transduction with CSIB-based virus followed by selection with 5 μg/ml blasticidin.

Cells were cultured in Dulbecco’s modified Eagle medium (DMEM) supplemented with 10% FCS at 37 °C and 5% CO_2_. Medium for Ace2-overexpressing cell lines contained 5 μg/ml blasticidin. All cell lines have been tested negative for contamination with mycoplasma and parental cell lines were obtained from the ATCC. Recombinant mouse-adapted SARS-CoV-2 MA virus was generated as described previously (GenBank Accession Number MT844088 (Dinnon et al., 2020). For virus titration, the caudal lobe of the right lung was homogenized in PBS, resulting homogenate was serial-diluted and inoculated onto confluent monolayers of Vero E6 cells, followed by agarose overlay. Plaques were visualized with overlay of Neutral Red dye on day 2 post infection (Dinnon et al., 2020).

### SARS-CoV-2/SARS-CoV-2 MA pseudotyped reporter virus

SARS-CoV-2 pseudotyped particles were produced by co-transfection of pNL4-3ΔEnv-nanoluc and pSARS-CoV-2-MA-S_trunc_ in 293T cell (Schmidt et al., 2020, Robbiani et al., 2020). For generation of SARS-CoV-2 MA pseudotyped particles, a plasmid expressing the mouse-adapted SARS-CoV-2 S (pSARS-CoV-2-MA-S_trunc_) was generated by introducing the Q498Y/P499T mutation into pSARS-CoV2-S_trunc_ and used for co-transfection with pNL4-3ΔEnv-nanoluc.

### SARS-CoV-2 MA pseudotyped virus neutralization assay

Four-fold serially diluted monoclonal antibodies were incubated with the SARS-CoV-2 MA pseudotyped virus for 1 hour at 37 °C. The mixture was subsequently incubated with HT1080_muAce2_ cells for 48 hours. Following incubation, cells were washed with PBS, lysed with Luciferase Cell Culture 5x reagent (Promega) and nanoluc luciferase activity in cell lysates was measured using the Nano-Glo Luciferase Assay System (Promega). Relative luminescence units obtained were normalized to those derived from cells infected with SARS-CoV-2 MA pseudovirus in the absence of monoclonal antibodies. The half-maximal and 90% inhibitory concentrations (IC50 and IC90) for monoclonal antibodies were determined using 4-parameter nonlinear regression (GraphPad Prism).

### Mouse studies and in vivo infections

All mouse studies were performed at the University of North Carolina (Animal Welfare Assurance #A3410-01) using protocols (19-168, 20-114) approved by the UNC Institutional Animal Care and Use Committee (IACUC). All animal work was approved by Institutional Animal Care and Use Committee at University of North Carolina at Chapel Hill according to guidelines outlined by the Association for the Assessment and Accreditation of Laboratory Animal Care and the U.S. Department of Agriculture. All work was performed with approved standard operating procedures and safety conditions for SARS-CoV-2. Our institutional BSL3 facilities have been designed to conform to the safety requirements recommended by Biosafety in Microbiological and Biomedical Laboratories (BMBL), the U.S. Department of Health and Human Services, the Public Health Service, the Centers for Disease Control and Prevention (CDC), and the National Institutes of Health (NIH). Laboratory safety plans have been submitted, and the facility has been approved for use by the UNC Department of Environmental Health and Safety (EHS) and the CDC. Twelve month old female BALB/c mice (Envigo, #047) were inoculated with the indicated concentration of antibody intraperitoneally 12 hours prior to infection. For infection, mice were anesthetized with a mixture of ketamine/xylazine and infected with 1×10^5^ PFU of SARS-CoV-2 MA in 50 μl PBS intranasally. Mice were daily monitored for body weight changes. At 2 days post infection, mice were euthanized, and lung tissue was harvested for virus titer analysis. Samples were stored at −80°C until homogenized and titered by plaque assay as described above.

## Acknowledgements

These studies were supported by George Mason University Fast Grants (T.P.S, M.C.N), a U19 grant from the National Institute of Allergy and Infectious Disease (1U19AI142759, Antiviral Drug Discovery and Development Center, R.S.B.) NIH grant P01-AI138398-S1 and 2U1 9AI111825 to M.C.N.; M.C.N. is a Howard Hughes Medical Institute Investigator. This project was supported by the North Carolina Policy Collaboratory at the University of North Carolina at Chapel Hill with funding from the North Carolina Coronavirus Relief Fund established and appropriated by the North Carolina General Assembly.

## Notes

### Competing Interest Statement

The authors have declared no competing interest.

